# Fast and Reliable: Evaluating Smartphone LiDAR App for Stem Diameter Measurement and Tree Mapping

**DOI:** 10.64898/2026.01.12.698980

**Authors:** Kirill Korznikov, Jiří Doležal, Pavel Fibich, Vojtech Lanta, Václav Bažant, Martin Aquino-Ramírez, Savanna Collins-Key, Jose Villanueva-Diaz, Jan Altman

## Abstract

Tree inventories require rapid, accurate measurements of stem diameter at breast height (DBH) and precise tree locations to support monitoring, planning, and informed decision-making. We evaluated a smartphone-based LiDAR app (SBLA), Forest Scanner, against (i) a diameter tape for DBH and (ii) a Vertex ultrasonic device for spatial coordinates. Across DBH of 725 trees, the LiDAR closely matched diameter tape measurements: discrepancies >5 cm occurred in 10.5% and > 10 cm in 3.5% of trees. Errors were concentrated in trees with smaller DBH, where occasional overestimation by SBLA arose from point-cloud misfitting. For medium and large trees, agreement was consistently high. Tree coordinates from SBLA and the ultrasonic device were broadly comparable at fine scales. Field efficiency was substantially improved: a 1,000 m^2^ plot with 70–80 trees required ∼2 hours using an ultrasonic device and diameter tape versus ∼20 minutes (one person) with SBLA, an ∼85–90% reduction in person-hours. Current limitations of SBLA are primarily software-related (stability, data handling, low-light performance). Overall, SBLA offers an efficient, auditable, and operationally relevant tool for tree inventories, with utility for rapidly updating DBH and spatial data used in management, planning, and asset databases.

## 1 INTRODUCTION

Diameter at breast height (DBH) is a core measurement in forest inventories that underpins key operational and ecological assessments in forestry (Guenther et al., 2024). Spatial distribution or mapping of trees, widely studied in forest ecology, also has important practical implications. In natural ecosystems, clustered distributions often reflect regeneration niches, dispersal limitation, or facilitative interactions, whereas regular or overdispersed patterns are typically associated with competition (Haase, 1995). In artificial forests and urban parklands, similar processes interact with human design and management decisions. For instance, planting schemes, soil sealing, or microclimatic gradients can either enhance clustering or promote uniform spacing to reduce competition and improve canopy distribution (McCoy et al., 2022). Spatial analyses of clustering and regularity can therefore serve as diagnostic tools, linking ecological understanding with forest management.

Thus, accurate and efficient methods for measuring DBH and tree positions are pivotal. Traditional forest inventory tools, such as measurement tapes and ultrasonic devices, are well established but can be labor-intensive, relatively slow, and error-prone in complex or dense understories. Recent advances in terrestrial Light Detection and Ranging tools, i.e., LiDAR (Holvoet et al., 2025), and the integration of LiDAR sensors into smartphones (Guenther et al., 2024; Gülci et al., 2023; Howie and De Stefano, 2024) offer the ability to capture tree dimensions and their spatial positions simultaneously via mobile applications, potentially reducing field time, minimizing transcription errors, and enabling frequent updates by small teams or even trained volunteers (Tatsumi et al., 2023). Our study evaluates the performance of smartphone-based LiDAR for forest inventory tasks (DBH and trees spatial mapping) and examines how method choice can influence downstream, ecological and practice-oriented interpretations.

Accordingly, we evaluate a smartphone-based LiDAR application (SBLA) (Tatsumi et al., 2023) for measuring DBH and tree spatial positions across a few forest types in Central Europe and North America. Specifically, we aimed to: (1) evaluate the accuracy of DBH measurements obtained from an SBLA compared with traditional measurements; (2) compare tree spatial coordinates measured by SBLA and an ultrasonic device; (3) test whether differences in measured tree positions alter spatial metrics.

## 2 METHODS

### 2.1 Study Area and Sampling Design

Tree measurements were conducted across ten sample plots: eight in Europe (the Czech Republic) and two in North America (Mexico and the USA). Czech plots were established in lowland temperate broadleaved forests composed mainly of *Acer campestre, Tilia cordata, Quercus robur*, and *Ulmus laevis* (plots B1–B6), and belonged to the nature reserve Soutok (48.72° N, 16.98° E) (Miklín and Čížek, 2014). Two other plots were established in the nature reserve Trebonsko (48.86° N, 14.80° E) (Kušová et al., 2008), where forests have a natural composition of coniferous trees *Picea abies, Pinus rotundata* and *Pinus sylvestris* (plots C1 and C2). The plot C3 was established in a mountain pine–oak woodland (*Pinus cembroides, Quercus* spp.) in central Mexico (20.40° N 98.49° W). The plot C4 was established in Texas (the USA), in a *Pinus taeda* forest stand under regular prescribed fire management (30.37° N 94.25° W).

Sampling was carried out in circular areas with a radius of 17.85 m (equal to the area of 1,000 m^2^) at all 10 plots. Within each plot, all trees with DBH ≥ 5 cm were measured. Tree density ranged from 22 to 86 trees per plot, reflecting substantial structural variation among sites. The total number of trees accounted for within the plots is 521, which is the overall number of trees used for comparison of spatial coordinates.

Additionally, we collected DBH data from two tropical lowland forest sites in Yucatán, Mexico (18.79° N, 88.49° W and 18.62° N, 90.56° W), two plots in each site. Across four circular plots, we measured a total of 204 trees, excluding palms. Thus, the total number of trees measured for the DBH comparison was 725.

### 2.2 Measurements

For all trees, we measured DBH at a height of 1.3 m above ground using a measurement tape. Tree spatial positions were obtained using an ultrasonic device Vertex Laser Geo 2 (Haglöf, 2021). We used ultrasonic measurement mode only, since laser mode was not possible due to the presence of understory plants between the device and the measuring trees. At the plot center, we mounted a base station on a monopod at a height of 1.3 m, operated by the first operator. The second operator carried a transponder mounted on a pole of the same height and sequentially positioned it at each tagged tree. The device measured horizontal distances and azimuths relative to the plot center, from which local Cartesian coordinates were calculated. During this process, the operator simultaneously measured tree DBH with a tape and identified tree species.

In parallel, we measured tree positions and DBH with an SBLA ForestScanner (v. 2.1) (Tatsumi et al., 2023) installed on an iPhone 15 Pro Max. The LiDAR sensor operates in the near-infrared spectrum with an effective range of 0.2–5 m. Each tree was scanned through the interface of SBLA by walking around the plot, during which all individual trees were marked within the app. Upon a tree selection, the application fitted a cylinder to the point cloud received from the stem surface at 1.3 m above ground to estimate DBH. Tree positions were simultaneously recorded relative to the scanning path and later exported as Cartesian coordinates.

### 2.3 Statistical Analyses

We compared DBHs derived from SBLA with tape measurements using linear regression and scatterplot visualization in R v.4.5.1 (R Core Team, 2025). Analyses were run for (a) all trees, and to account for morphological and biogeographical differences, separately for (b) broadleaf temperate trees, (c) coniferous temperate trees, and (d) broadleaf tropical trees. To assess systematic bias, we computed the arithmetic difference between SBLA and tape measurements.

To compare differences in spatial positions of trees from the ultrasonic device and the SBLA, we calculated two plot-level summary metrics: mean pairwise distance (MPD) and mean nearest-neighbor distance (MNND). Both metrics capture complementary aspects of stand structure. MNND reflects fine-scale spacing and local crowding, which influence tree growth, management needs, and risks such as branch conflict. MPD, in contrast, summarizes broader stand configuration and overall tree distribution within a plot. Together, the metrics provide a straightforward and interpretable way to assess whether measurement differences between tools are large enough to affect ecological or management-relevant assessments of spatial structure (Szmyt, 2014).

Also, for spatial pattern analysis, we applied Ripley’s *K*-function and the mark correlation function (*K*_*mm*_) with DBH as the mark. *K*-function is widely used in forest ecology because it characterizes spatial structure across multiple distances, enabling detection of spatial clustering or regularity relative to some null/neutral distribution, i.e. complete spatial randomness where individuals occur within the plot in random places (Haase, 1995; Velázquez et al., 2016). The mark correlation function assessed size-related associations by testing if lower or higher than mean marks are spatially associated relative to some null distribution, i.e. permutation of marks. For each plot, we generated 499 null distributions by: (1) complete spatial randomness for Ripley’s *K* function, and (2) randomly permuting DBH values over fixed tree locations for the mark correlation function. Significance was evaluated using the Diggle–Cressie–Loosmore–Ford test up to 5 m, with isotropic edge correction applied throughout. All analyses were conducted in R v.4.5.1 (R Core Team, 2025). using the *spatstat* package (v. 3.0-8) (Baddeley and Turner, 2005). Code is provided in the Supplementary Materials.

## 3 RESULTS

### 3.1 DBH measurements

DBH values obtained from the SBLA were strongly correlated with those measured using a tape (Fig. 1a). Discrepancies greater than 5 cm occurred in 10.5% of cases, and discrepancies exceeding 10 cm were observed in only 3.5%. Like the overall datasets, tree groups also showed a highly significant and positive linear relationship between SBLA- and diameter tape-measured DBH values (Fig. 1b–d).

**Figure 1.**
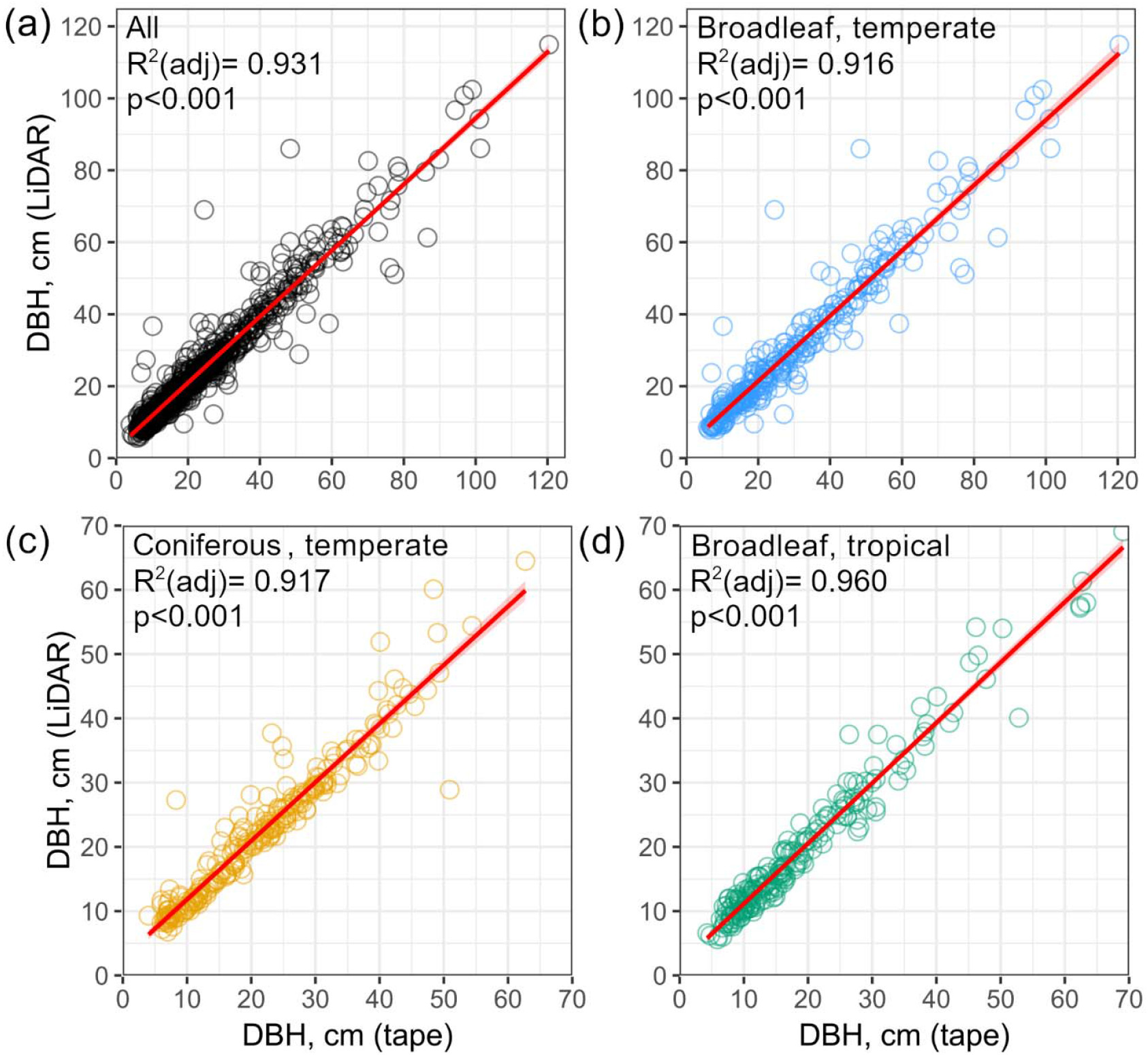
Comparison of smartphone-based LiDAR application and tape-measured DBH across: (a) all trees, (b) broadleaf temperate trees, (c) coniferous trees, and (d) broadleaf tropical trees; linear regression lines with 95% confidence intervals are shown.

When examining deviations directly, measurement errors were most pronounced in small tree stems, whereas larger trees showed close agreement between methods (Fig. 2). The magnitude of discrepancies declined with increasing DBH, indicating that the SBLA provides more consistent and reliable estimates for medium and large trees.

**Figure 2.**
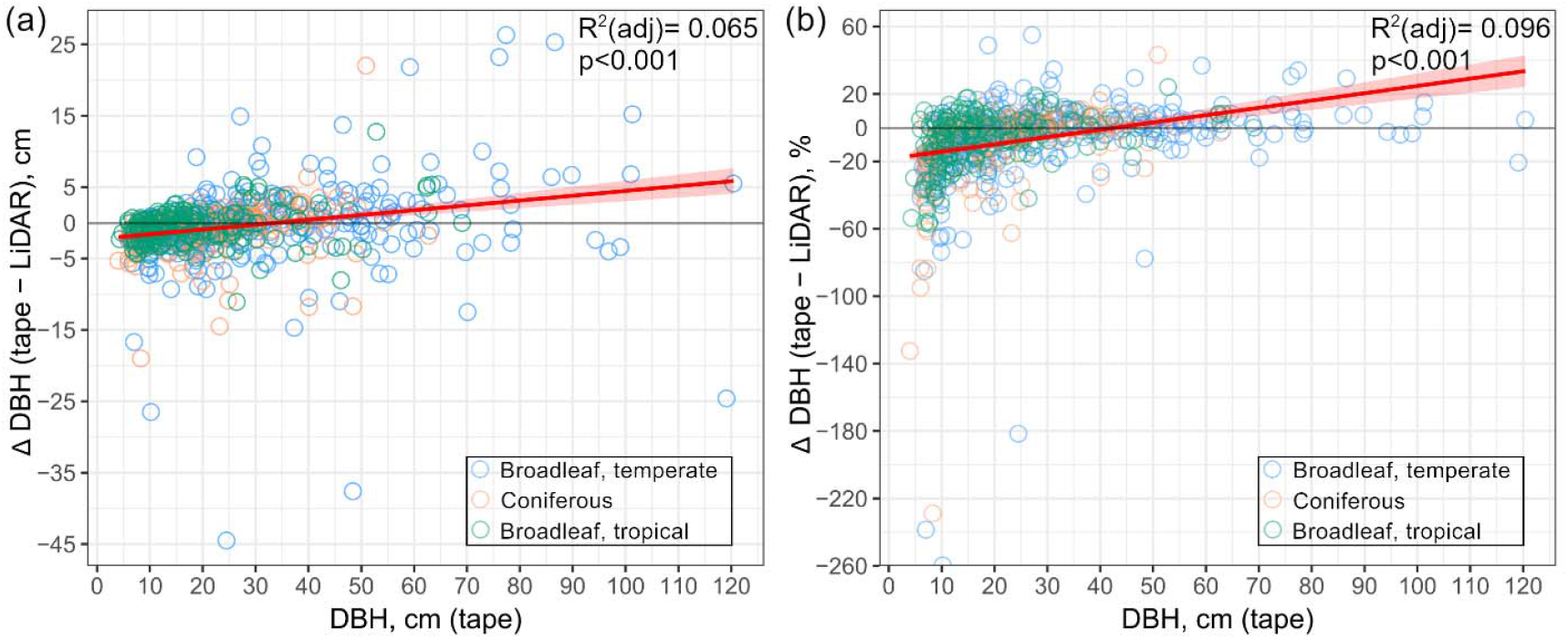
Differences between smartphone-based LiDAR application and tape-measured DBHs are shown in (a) absolute values and (b) as a percentage of the tape measurement; positive values indicate smartphone-based LiDAR application underestimation.

### 3.2 Distances among trees

Statistically significant differences in MNND were detected in two sample plots (Fig. 3a), both showing higher values by the ultrasonic device. At the MPD scale, differences between methods became more pronounced (Fig. 3b). We found significant deviations in nine plots, seven of which showed higher MPD values from the ultrasonic device and two in which SBLA values were higher. One sample plot showed no significant difference. The largest discrepancy occurred at plot B6, where ultrasonic device-based MPD exceeded SBLA estimates by nearly 7 m.

**Figure 3.**
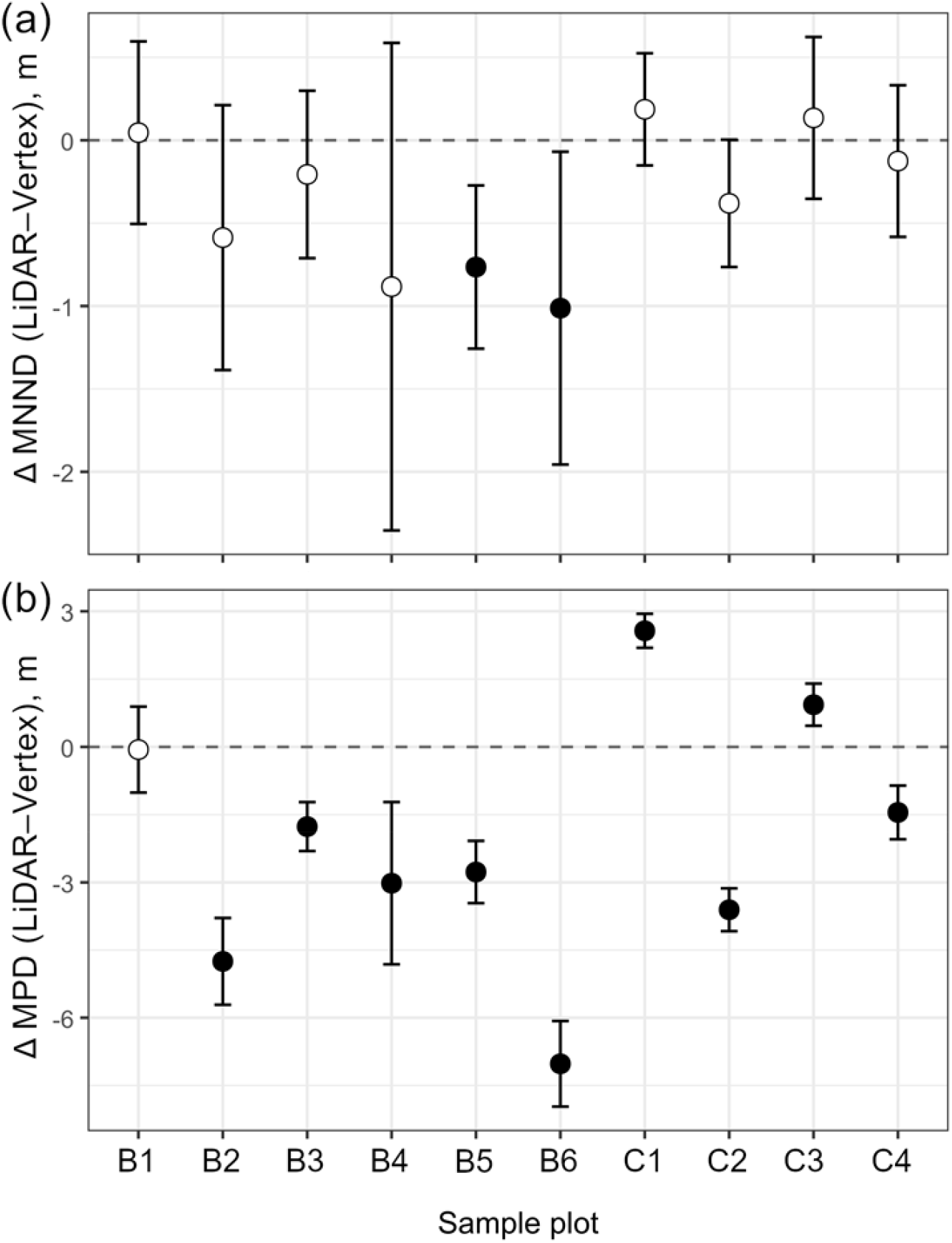
Differences in (a) mean nearest-neighbor distance (MNND) and (b) mean pairwise distance (MPD) between smartphone-based LiDAR application and ultrasonic device-derived across sample plots; values on Y-axes represent differences with 95% confidence intervals, the dashed line indicates no difference; filled symbols denote statistically significant differences by the paired t-test (p < 0.05), whiskers show 95% confidence intervals; plot prefixes: B – broadleaf forest stands, C – conifer or coniferous-dominated forest stands.

### 3.3 Spatial structure

Spatial pattern analyses using Ripley’s *K*-function and the mark correlation function showed broadly consistent results between SBLA and ultrasonic device-derived tree positions. In most plots, both methods indicated spatial clustering of trees and weak or absent DBH-based associations. The discrepancies between methods occurred at only one sample plot (C2), where ultrasonic device data revealed significant tree clustering (Fig. 4a) and DBH-related associations (Fig. 4b), whereas SBLA data did not. Across all other plots, both approaches produced mostly concordant outcomes, with observed values remaining within the simulation envelopes of the null models (Fig. S1).

**Figure 4.**
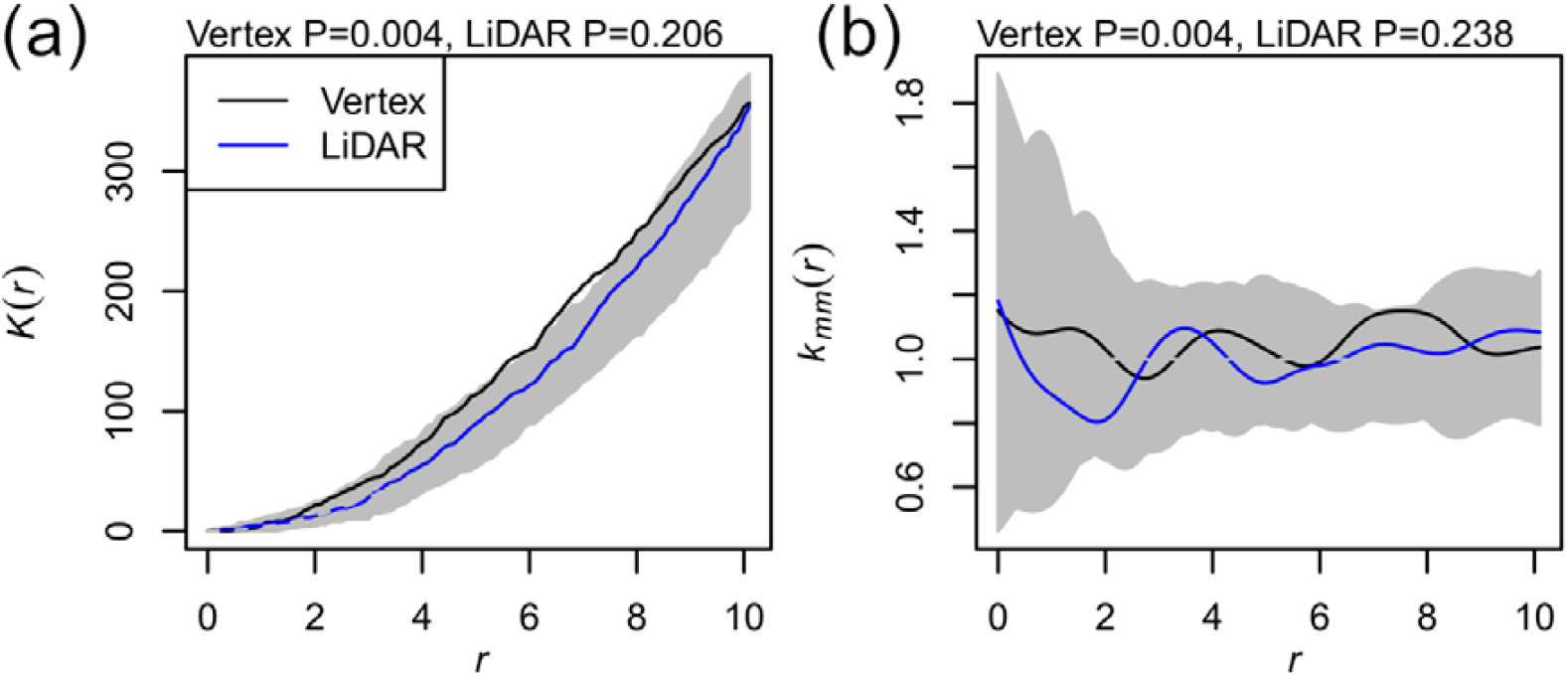
Spatial pattern analysis of tree positions and DBH values on the plot C2(n=79) based on smartphone-based LiDAR application and ultrasonic Vertex device measurements, (a) Ripley’s K-function (*K*) and (b) mark correlation function (*K*_*mm*_); grey area in all panels indicates 95% confidence envelope generated from 499 null distributions; (a) shows spatial associations whether tree positions are random, clustered, or regularly spaced at increasing spatial scales; lines above the grey envelope indicate clustering (trees are closer together than if they would be distributed randomly), lines below the envelope indicate regularity (trees are more evenly spaced than if they would be distributed randomly), lines within the envelope indicate a pattern consistent with the random distribution. (b) shows whether tree DBH values are spatially associated; values higher or lower than 1 signify that, within a given spatial scale, individuals tend to have higher or lower DBH than the plot-wide mean value; values near 1 indicate no size-related spatial association; P-values <0.05 show significance of deviation from the null distribution by Diggle–Cressie–Loosmore–Ford tests for distances up to 5m.

## 4 DISCUSSION

### 4.1 DBH

SBLA provided DBH estimates that generally corresponded well with tape measurements. In our study, most extreme outliers occurred among small trees, where SBLA occasionally overestimated stem size due to misfitting of the trunk contour, when point clouds were sparse or obscured. One stem with the measured by tape DBH 24.5 cm was recorded by SBLA as 62.7 cm, highlighting that such errors can also occur outside the smallest size class, though rarely acros studied trees.

Recent studies using smartphone-based LiDAR sensors also report strong DBH correlation with traditional methods (R^2^ >0.88) but also note decreasing accuracy for small diameters and irregular stem forms (Guenther et al., 2024; Gülci et al., 2023; Howie and De Stefano, 2024). Thus, SBLA- and tape-based DBH measurements were generally consistent, but SBLA did not universally outperform diameter tapes. Its main advantage emerged for larger stems, whereas small trees were often overestimated due to sparse or incomplete point clouds. In addition, buttressed stems, irregular growth forms, or the presence of epiphytic species attached to tree trunks can complicate stem geometry and lead to substantial errors in DBH estimation.

### 4.2 Spatial position

Tree spatial positions derived from SBLA and the ultrasonic device were generally consistent, although the ultrasonic device tended to produce slightly larger inter-tree distances. However, in most cases, we did not find discrepancies in spatial patterns revealed by Ripley’s *K*-function and the mark correlation function. Divergent spatial interpretations occurred in only one sample plot, where SBLA data indicated significant tree clustering while ultrasonic data suggested a random distribution. Such a case highlights that the choice of field measurement technique can directly affect conclusions about forest stand structure and spatial organization.

Ultrasonic devices tend to overestimate inter-tree distances relative to SBLA, a bias likely caused by cumulative angular errors and by measuring to the trunk surface instead of the stem centre. The assumption that these discrepancies would propagate into ecological analyses was only partly supported. In most plots, both methods produced similar tree spatial patterns.

### 4.3 Practical outcomes

Both measuring approaches, an ultrasonic device combined with measuring tapes and SBLA, offer distinct advantages and limitations that determine their suitability for tree measurements. Our findings and field experience indicate that tape measurements of DBH may be inaccurate, whether from rushing in the field, misreading or misplacing the tape, or transcription mistakes during data entry. In contrast, SBLAs offer the advantage of a permanent digital record, providing an additional layer of quality control that traditional methods cannot offer. Regarding spatial positions, the ultrasonic device is prone to line-of-sight disruptions, whereas the SBLA is constrained by its effective range (≤ 5 m), occasional software instability, and high smartphone battery consumption. Neither approach is error-free, and their performance depends strongly on stand structure, understory density, and the experience of operators.

Several factors may contribute to systematic error in ultrasonic measurements. Small angular deviations during device rotation, particularly in dense stands, can accumulate into measurable positional shifts. Furthermore, because the transponder is mounted on the outer trunk surface, recorded distances reflect stem edges rather than stem centres, introducing a bias that increases with tree size.

SBLA ForestScanner (Tatsumi et al., 2023) is not free of limitations either. During fieldwork, some trees appeared displaced on the device screen, reflecting memory constraints, transient data misalignment, or errors in automatic tree stem recognition. Reduced performance was also observed under low-light conditions, where the ultrasonic device continued to operate reliably. Also, the precision of SBLA in estimating true stem centres remains uncertain and warrants further validation. Preliminary checks suggest improvement relative to the ultrasonic device, but misfitting point clouds for banding stems can still introduce distortions.

Overall, our results suggest that SBLAs can substitute for tape measurements for medium and large trees, but caution is needed, especially for small stems, where overestimation is more likely. Both techniques face challenges in accurately capturing multistem trees, where defining a clear geometric centre is difficult. Users must therefore be prepared to measure and correctly interpret “challenge” tree forms, as they represent an integral component of natural forests. Importantly, rather than viewing tape as the “true” baseline in our study, both approaches should be seen as complementary, with LiDAR adding transparency and verifiability to the measurement process.

The clear strength of SBLAs lies in their efficiency. In this study, a 1,000 m^2^ plot containing 70–80 trees required approximately two hours for two operators using the ultrasonic device and measuring tape, but only about 20 minutes when measured by a single operator with a smartphone. This represents an 85–90% reduction in person-hours, a substantial saving for large-scale or long-term forest monitoring. Importantly, SBLA eliminates the space for follow-up transcription errors as it directly records all measurements. In contrast, the ultrasonic device records information about spatial positions, but DBH (and other information) needs to be digitalized after fieldwork. In addition, archived scans made by SBLA can be easily revisited for validation or reanalysis (Fig. S2).

Despite the benefits, SBLA is not flawless. Occasional problems with tree recognition, memory limitations, or reduced accuracy under low-light conditions affected measurements. These issues stem from the current generation of applications rather than the LiDAR sensor itself. Future improvements in app stability, interface design, and automated error detection are likely to enhance data quality and reliability. Moreover, the development of dedicated applications could further expand the potential of SBLAs by integrating standardized measurement protocols, automated DBH calibration, and cloud-based data management.

Compared with larger terrestrial LiDAR systems (Oveland et al., 2017), SBLA is far more portable and accessible, but sacrifices range, resolution, and automation. Its scanning limit remains a critical constraint, reducing reliability for very large stems and complicating data acquisition in dense stands. Nevertheless, the combination of portability, rapid data collection, and direct digital archiving makes SBLA a valuable complement to conventional field methods, particularly for inventory projects constrained by time, personnel, or funding. In addition, its ease of use and compatibility with mobile data platforms make it an ideal tool for forestry applications, where accurate and up-to-date tree information supports monitoring, planning, and the development of spatial databases for green infrastructure management.

## CONCLUSION

SBLAs offer a promising and practical complement to traditional tree measurement approaches. Our results demonstrate that SBLA already provides an efficient and sufficiently accurate alternative to tape DBH measurements and ultrasonic device-based trees position mapping, substantially reducing field effort and personnel requirements. However, app instability, high battery consumption, and limited data management functions remain key challenges that still face limitations, hindering immediate integration into standard tree inventory protocols. Importantly, the constraints stem from software implementation rather than from the LiDAR technology itself and can likely be resolved through further updates or the development of new, purpose-built applications. Such tools should integrate enhanced data storage and synchronization, automated quality control, improved stem recognition and calibration, real-time error detection, and seamless links to geospatial and inventory databases – features essential for meeting the diverse requirements of both forestry monitoring and ecological research.

## Acknowledgment

We thank all colleagues who contributed, in various capacities, to the selection of study sites and to the organization and execution of field measurements in the Czech Republic, Mexico, and the United States, as well as those who assisted with the processing of field data.

## CRediT authorship contribution statement

**Kirill Korznikov**: Conceptualization, Data Curation, Investigation, Formal Analysis, Methodology, Visualization, Writing – Original Draft Preparation, Writing – Review & Editing. **Ji**ř**í Doležal**: Conceptualization, Investigation, Validation, Writing – Review & Editing. **Pavel Fibich**: Formal Analysis, Software, Validation, Visualization, Writing – Review & Editing. **Vojtech Lanta**: Investigation, Writing – Review & Editing. **Václav Bažant**: Investigation, Writing – Review & Editing. **Martin Aquino-Ramírez**: Investigation, Writing – Review & Editing. **Savanna Collins-Key:** Investigation, Writing – Review & Editing. **Jose Villanueva-Diaz**: Investigation, Writing – Review & Editing. **Jan Altman**: Conceptualization, Funding Acquisition, Methodology, Validation, Writing – Original Draft Preparation, Writing – Review & Editing.

## Supplementary figures

**Figure S1.**
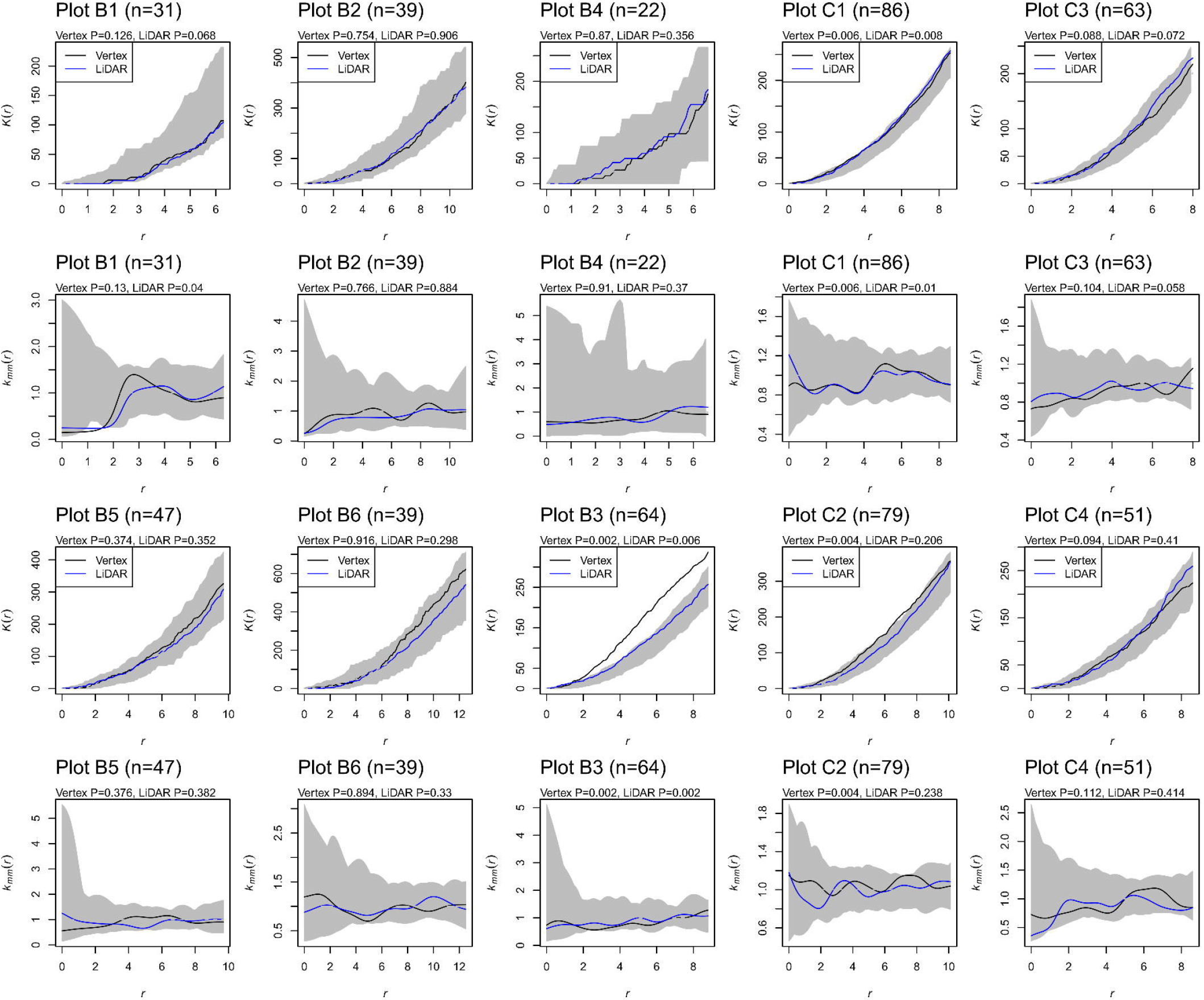
Spatial pattern analysis of tree positions and DBH values based on smartphone-based LiDAR application and ultrasonic Vertex device measurements. Grey area in all panels indicates 95% confidence envelope generated from 499 complete spatial randomness simulations (null model). The upper graph in each box shows spatial clustering, whether tree positions are random, clustered, or regularly spaced at increasing spatial scales; lines above the grey envelope indicate clustering (trees are closer together than random), lines below the envelope indicate regularity (trees are more evenly spaced than random), and lines within the envelope indicate a pattern consistent with randomness. The lower panel shows whether tree DBH values are spatially associated; values > 1 indicate trees of similar DBH are closer than expected (size-based clustering); values < 1 indicate small and large trees are more intermixed than random; values near 1 indicate no size-related spatial association. P-values <0.05 show significance of deviation from the null model.

**Figure S2.** Spatial comparison of forest inventory maps derived from two independent measurements by the smartphone-based LiDAR app ForestScanner at the sample plot B3 in the Czech Republic; green circles indicate *Quercus robur* trees, violet circles represent *Acer campestre* trees, circle size is proportional to measured DBH; differences in coordinates reflect inaccuracies in the initial smartphone GPS positioning.

