## Supplementary figures and images for "Fast and Reliable: Evaluating Smartphone LiDAR App for Stem Diameter Measurement and Tree Mapping"

### Figure S2

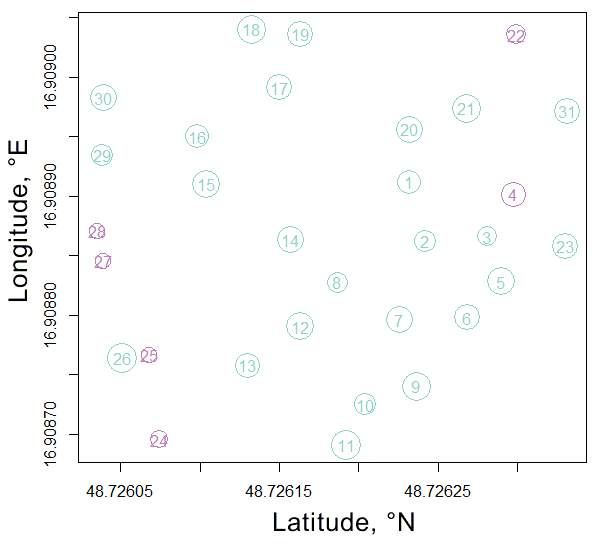
